# Wide Diversity of Recombinant Noroviruses Circulating in Spain, 2016 to 2020

**DOI:** 10.1101/2021.12.02.471049

**Authors:** Noemi Navarro-Lleó, Cristina Santiso-Bellón, Susana Vila-Vicent, Noelia Carmona-Vicente, Roberto Gozalbo-Rovira, Jesús Rodríguez-Díaz, Javier Buesa

## Abstract

Noroviruses are the leading cause of sporadic cases and outbreaks of viral gastroenteritis. For more than 20 years most norovirus infections have been caused by the pandemic genotype GII.4, yet recent studies have reported the emergence of recombinant strains in many countries. In the present study, 4,950 stool samples collected between January 2016 and April 2020 in Valencia (Spain) from patients with acute gastroenteritis were analyzed to investigate the etiological agent. Norovirus was the most frequently detected enteric virus with a positive rate of 9.5% (471/4,950). Among 224 norovirus strains characterized, 175 belonged to genogroup GII and 49 to genogroup GI. Using dual genotyping based on sequencing the ORF1/ORF2 junction region we detected 25 different capsid-polymerase type associations. The most common GII capsid genotype was GII.4 Sydney 2012, followed by GII.2, GII.3, GII.6 and GII.17. A high prevalence of recombinant strains (90.4%) was observed among GII infections between 2018 and 2020. GII.4 Sydney[P16] was the predominant genotype from 2019 to 2020. In addition, GII.P16 polymerase was found harboring within six different capsid genes. A new subcluster of GII.4 Sydney associated with the P31 polymerase was identified by phylogenetic analysis. GI.4 and GI.3 were the predominant genotypes in genogroup GI, in which recombinant strains were also found, such as GI.3[P10], GI.3[P13] and GI.5[P4]. Interestingly, the GI.3[P10] strain could represent a new capsid genotype. This study shows the extensive diversity of recombinant noroviruses circulating in Spain and highlights the role of recombination events in the spread of noroviruses.

## INTRODUCTION

Norovirus (NoV) infections are considered a primary cause of acute gastroenteritis (AGE) in all age groups worldwide (1, 2) and lead to a major health problem. In developed countries norovirus outbreaks bring a high economic burden, estimated at over $64 billion annually (3). The settings most affected are hospitals (4), restaurants (5), cruise ships (6), schools (7), and nursing homes (8). In countries where rotavirus vaccination has been implemented, noroviruses have become a major cause of gastroenteritis in children (9-11). These viruses are highly contagious and are transmitted person to person and by contaminated food or water (12). It has been estimated that noroviruses cause between 70,000 and 200,000 deaths annually in developing countries (2, 13).

The genus *Norovirus* belongs to the *Caliciviridae* family, which are non-enveloped viruses with a single-stranded positive-sense RNA genome of approximately 7.5 kb in length, organized into three open reading frames (ORFs). ORF1 encodes a polyprotein that is post-translationally cleaved into six non-structural viral proteins including the RNA-dependent RNA polymerase (RdRp). ORF2 and ORF3 encode the major (VP1) and minor (VP2) capsid proteins, respectively (14). VP1 is organized into the shell (S) and protruding (P) domains (13). The P domain can be further divided into P1 and P2 subdomains. The P1 subdomain forms the anchoring portion of the P dimer, connecting it to the S domain, while the highly variable P2 subdomain is the most surface-exposed region of the norovirus capsid. The P2 subdomain acts as the target for neutralizing antibodies and is the receptor binding site for histo-blood group antigens (HBGAs) (15).

Based on phylogenetic analyses of the complete VP1 amino acid sequences, norovirus have been classified into 10 distinct genogroups (GI-GX), which are subdivided into 49 different genotypes (16). Of these, GI, GII and GIV strains cause illness in humans, and the majority of diseases are correlated with GI and GII infections.

Norovirus diversity arises through evolution mechanisms such as recombination or accumulation of mutations (17). Recombination often occurs at the ORF1/ORF2 junction, leading to new combinations of capsid and RdRp genotypes which contribute to increase genetic diversity (17-19). As a result, these new recombinant strains might have increased fitness, pathogenicity and/or transmissibility compared to their ancestor strains (20-22). Furthermore, the same capsid harbors different RdRp genotypes, which may offer an advantage by changing the efficiency of virus replication (23). Despite the high genetic diversity of noroviruses, during recent decades a single genotype (GII.4) has been the most prevalent in humans worldwide (24). This virus is highly versatile and new variants emerge rapidly due to antigenic drift in the VP1 and to genetic recombination between circulating norovirus strains (19, 25).

To understand the epidemiology and the genotypic trends of evolving noroviruses, we studied and characterized the molecular epidemiology of norovirus strains causing sporadic cases or outbreaks of AGE in the Valencian Community (Spain) from January 2016 to April 2020.

## MATERIALS AND METHODS

### Sample collection

Presence of noroviruses was investigated in specimens from patients with sporadic gastroenteritis attended at primary care clinics and at the emergency department of the Hospital Clínico Universitario of Valencia. A total of 4,950 stool samples were tested by conventional RT-PCR and/or RT-qPCR between January 2016 and April 2020. The study was approved by the Clinical Research Ethics Committee of the Hospital Clínico Universitario of Valencia. Additionally, samples from norovirus gastroenteritis outbreaks occurring at different Spanish locations (Alicante, Castellón, Cádiz, San Sebastián and Palma de Mallorca) were also analyzed anonymously. All specimens were stored at 4ºC until further molecular analysis.

### RNA extraction and reverse transcription

Viral RNA was extracted from 10% fecal suspensions in PBS with TRIzol® reagent (Invitrogen) (26). The reverse transcription reaction was performed with random primers using SuperScript® III reverse transcriptase (Invitrogen).

### Norovirus detection

From January 2016 to May 2018 viral detection was carried out by conventional PCR, amplifying the polymerase (JV12/JV13 primers) and/or the capsid gene (G1SKF/G1SKR and G2SKF/G2SKR for GI and GII, respectively) (27, 28). Amplicons were analyzed by electrophoresis in 1.5% agarose gels with RedSafe staining (RedSafe™ Nucleic Acid Staining Solution, iNtRON Biotechnology). After June 2018, presence of noroviruses and other enteric viruses in stool samples was determined using the BD Max™ Enteric Viral Panel on the BD Max™ system (Becton Dickinson).

### Detection and analysis of recombinant strains

The overlapping ORF1/ORF2 junction region was sequenced to identify recombinant norovirus strains. In brief, RT-PCR was performed with a combination of previously described oligonucleotides: primers MON 432 (TGG ACI CGY GGI CCY AAY CA) and G1SKR (CCA ACC CAR CCA TTR TAC A) for genogroup I; primers MON 431 (TGG ACI AGR GGI CCY AAY CA) and G2SKR (CCR CCN GCA TRH CCR TTR TAC AT) for genogroup II (28, 29). The RT-PCR was performed using Qiagen One-Step RT-PCR kit (Qiagen) master mix, with 20 U of RNase inhibitor (Biotools) and the following thermal cycling conditions: 30 min at 50ºC, 15 min at 95ºC, and 40 cycles of 95ºC for 30 sec, 50ºC for 30 sec, and 72ºC for 45 sec, followed by 7 min at 72ºC.

To verify the recombination event, the ORF1/ORF2 junction fragments constructed by PCR as described above were analyzed along with reference strains obtained from GenBank by using SimPlot software v.3.5.1. The SimPlot analysis was performed by setting the window width and step size to 200 bp and 20 bp, respectively.

### Sequencing and genotyping method

The PCR products were purified by using NZYGelpure kit (Nzytech) and sequenced upstream and downstream. Sequence quality was checked with BioEdit software v7.0.0 (30). The viral genogroup, genotype and variant were assigned by using the Norovirus typing tool (http://www.rivm.nl/mpf/norovirus/typingtool) and the Human Calicivirus typing tool (http://norovirus.ng.philab.cdc.gov/). The strains were named following the new proposed dual-typing designation indicating the genotype of the capsid followed by the polymerase type in brackets (16).

### Data and phylogenetic analyses

Temporal distribution of norovirus infections per quarter and year was statistically analyzed with GraphPad Prism software, version 9.3.0. Pearson’s chi-squared test was performed to evaluate monthly quarters’ sporadic cases for which the proportion each year differed from a hypothetical 25%.

Phylogenetic trees were built with the sequences of this study and reference sequences obtained from GenBank. Sequences were aligned with Clustal X 2.0 (31) and equated with GeneDoc 2.7.000 (32). Finally, phylogenetic analyses were performed using MEGA7 (Molecular Evolutionary Genetics Analysis v7.0) (33). The MEGA maximum likelihood model selection tool was used to determine the best model for branch support using the Bayesian Information Criterion for analysis. Furthermore, the best model suggested by the program was used to calculate the level of nucleotide sequence identity between the sequences studied. The evolutionary history was inferred by the Maximum Likelihood method (34) or Neighbor joining method (35) using a bootstrap test of 1000 replicates to evaluate tree reliability.

### Accession numbers

Norovirus sequences derived in this study were deposited in the GenBank under accession numbers MN918435-MN918437, MN854082-MN854085, MT491997-MT492003, MT492038-MT492049, MT492061-MT492069, MT495616-MT495630, MT495732-MT495748, MT501813-MT501864, MT908120-MT908122 and MT908849-MT908867.

## RESULTS

### Norovirus detection and typing

In total, 9.5% (471/4,950) stool specimens tested norovirus positive by conventional RT-PCR and/or RT-qPCR. From these, 264 viral strains could be further amplified for sequencing, genogrouping and genotyping, which was eventually achieved with 224 strains (84.8%). Their polymerase and/or capsid genotypes were determined, 49 (21.9%) strains belonging to genogroup I (GI) and 175 (78.1%) strains to genogroup II (GII).

### Genetic diversity of noroviruses

A wide range of norovirus genotypes was detected throughout the 5-year study period (Table I). Six capsid genotypes were identified within genogroup GI (GI.1, GI.3-GI.7) and 14 capsid genotypes in GII (GII.1-GII.7, GII.10, GII.12-GII.14, GII.17, GII.20 and GII.21). The most prevalent GI genotype was GI.4 (13/40, 32.5%), which was involved in four different outbreaks occurring in March 2018 and 2019 (Fig.1A). Strains showing the capsid gene GI.3 represented the second most frequent genotype (10/40, 25%), and it was found mainly from 2018 onwards (Fig.1A). In genogroup II, GII.4 viruses caused 47.9% (58/121) of AGE cases. The GII.4 Sydney variant represented 50.8% (31/61) of norovirus strains detected in 2019 (Fig.1A). There were 17 GII.4 isolates that could not be assigned to a specific variant which were associated with P4 New Orleans or P31 polymerases (Fig.1A). Other frequently identified genotypes along this study were GII.2, GII.6 and GII.17 at the same ratio (12/121, 9.9%) and GII.3 (7/121, 5.8%). It is noteworthy that GII.17 was detected late in 2018 (Fig. 1A). The temporal distribution of polymerase genotypes showed that GII[P17] and GII[P4] predominated from 2016 to 2018 with the same prevalence (19/84, 22.6%) and were replaced by GII[P16] (42/72, 58.3%) from 2019 onwards (Fig.1B). Furthermore, GII[P4 New Orleans] and GII[P31] were widely detected in the same proportion (12.2%) throughout the study period (Fig.1B).

**Table 1.**
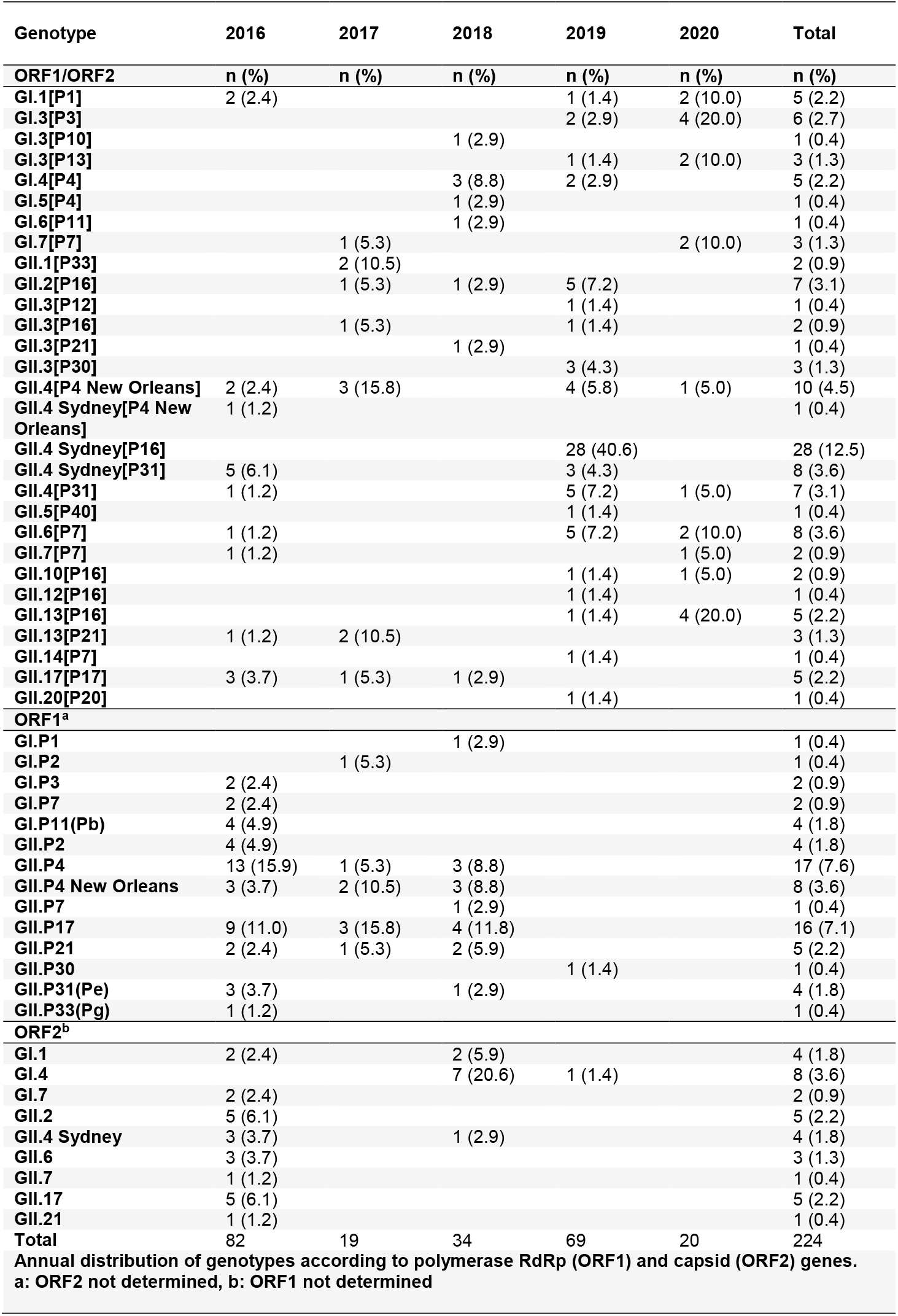
Norovirus genotypes producing sporadic cases or outbreaks of gastroenteritis in Spain (Valencian Community), from January 2016 to April 2020

**Fig. 1.**
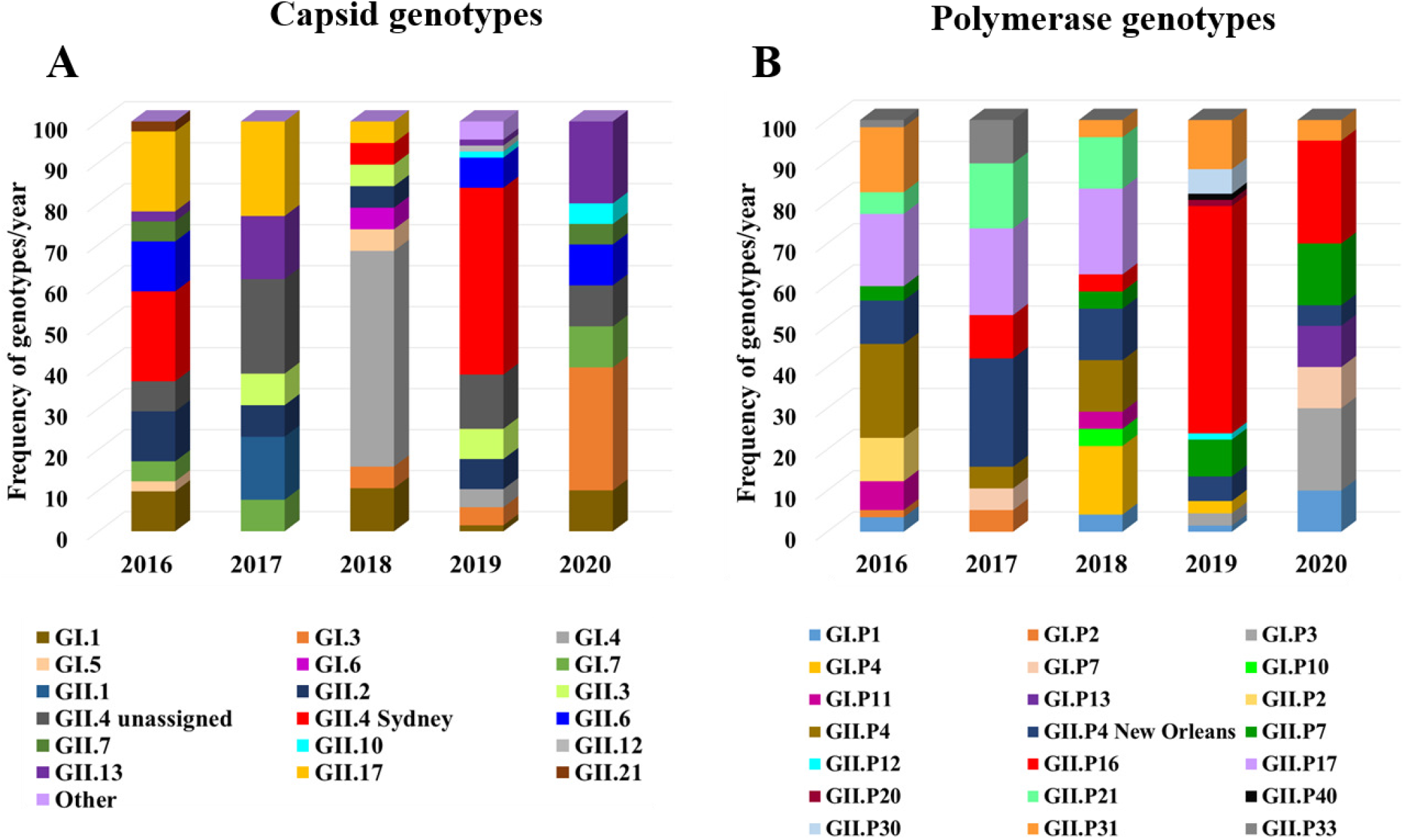
Distribution by year of GI and GII norovirus genotypes of sporadic cases or outbreaks in the Valencian Community, Spain, between January 2016 and April 2020. A) Capsid genotypes. Others include GII.5, GII.14 and GII.20. B) Polymerase genotypes.

### Temporal distribution of noroviruses

Of the 264 norovirus cases, 83 (31.4%) were detected in 2016, 40 (15.2%) in 2017, 46 (17.4%) in 2018, 75 (28.4%) in 2019 and 20 (7.6%) between January and April 2020. The monthly distribution of acute gastroenteritis due to noroviruses from 2016 to 2020 showed clear seasonality (Fig. 2). The prevalence of norovirus infections was significantly higher during the first quarter of the year in 2016 and 2018, but during the last quarter in 2019 (*P* < 0.05). Peaks of cumulative norovirus cases were observed in March and October (2016 and 2019) or November (2017 and 2018) while in contrast few sporadic norovirus cases were registered in July and August.

**Fig. 2.**
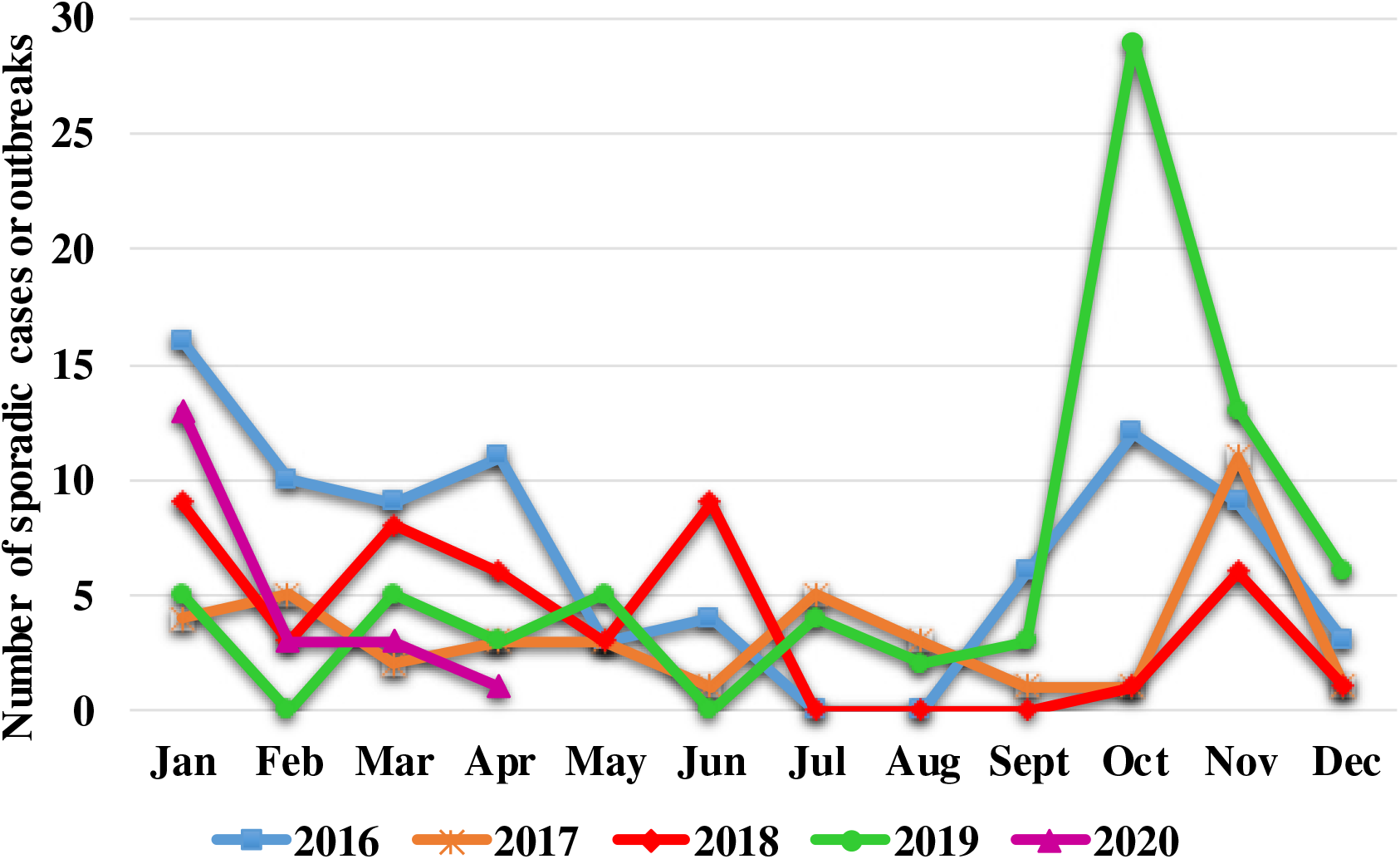
Distribution by month of norovirus infections detected from 2016 to 2020 in the Valencian Community, Spain.

### Recombinant norovirus strains

From January 2016 to May 2018 six recombinant norovirus genotypes were detected, including GII.1[P33], GII.2[P16], GII.3[P16], GII.4 Sydney[P31], GII.6[P7] and GII.13[P21]. From June 2018 onwards all norovirus positive samples by BD Max™ were routinely analyzed by sequencing the ORF1/ORF2 junction to detect recombinant genotypes, yielding polymerase and capsid sequences for 90 norovirus strains detected between June 2018 and April 2020. A remarkable number of recombinant strains belonged to genogroup II (90.4%, 66/73). Genotypes GII.4 Sydney[P16] (38.4%, 28/73), GII.6[P7] (7/73, 9.6%), GII.4[P31] (6/73, 8.2%) and GII.2[P16] (6/73, 8.2%) were found during the 2019– 2020 season (Fig. 3C). Several capsid genotypes were associated with more than one polymerase type, including GI.3, GII.3 and GII.4 Sydney. Viruses harboring the GII[P16] polymerase were detected in 59% (43/73) of AGE cases. This GII[P16] polymerase was found among cases involving GII.2, GII.3, GII.4 Sydney 2012, GII.10, GII.12 and GII.13 viruses. Furthermore, noroviruses combined with the GII[P31] (Pe) polymerase were associated with the GII.4 Sydney variant, although also with GII.4 unassigned strains. Viruses with GII[P12], GII[P21] and GII[P30] (Pc) polymerases were associated with GII.3. We also detected recombinant strains belonging to genogroup GI, such as GI.3[P10], GI.3[P13] and GI.5[P4].

**Fig. 3.**
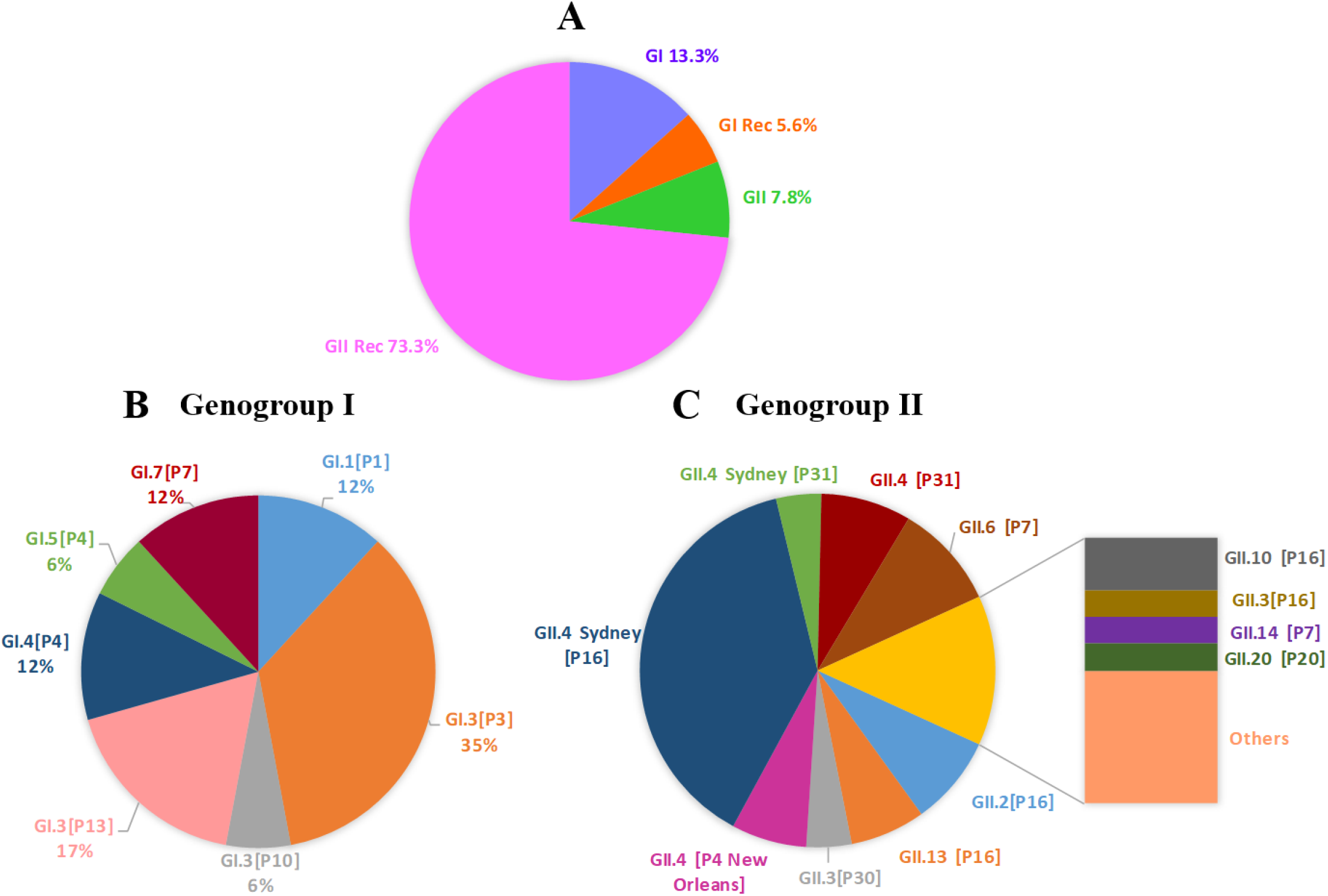
Diversity of norovirus genotypes from June 2018 to April 2020 in the Valencian Community, Spain. A) Genogroup I, genogroup II and recombinant strains. B) Distribution of genogroup I. C) Distribution of genogroup II strains. Others include GII.3[P12], GII.3[P21], GII.5[P40], GII.7[P7] and GII.12[P16].

#### Genetic analysis of norovirus genogroup GI

##### Phylogenetic analysis

ORF1/ORF2 junction sequence analysis of 19 GI norovirus strains revealed a division into two main clusters, one including GI.1[P1], GI.3[P3], GI.3[P10], GI.3[P13] and GI.4[P4] genotypes (cluster A) and the other one including only the GI.7[P7] genotype (cluster B) (Fig. 4). Cluster A is subdivided into two well-defined subclusters, one of them made up of genotypes associated with the GI.3 capsid. Regarding GI.3[P3] strains, two subgroups were observed, one close to a recent 2019 reference strain (MT031988) with a high nucleotide identity (98.7%), and another clustering with an older 2007 reference strain (NC_039897) with a 96.8% identity. The recombinant GI.3 “untypeable” [P10] strain detected in this study showed a low degree of nucleotide identity (87%) with reference GI.3[P10] strains (U04469 and MN421587). In addition, in the GI.3[P13] subcluster, two subgroups were observed, one grouped with an old 2010 variant (JQ911594) with a high identity (96.8%), and the second with 99.8% sequence identity with a recent 2019 variant (MN922742). GI.1[P1] and GI.4[P4] strains formed the second subcluster within cluster A, and grouped with more recent reference strains (MK956177 and MK956173) for GI.4[P4] and GI.1[P1] genotypes, respectively. Finally, cluster B includes only GI.7[P7] strains, which grouped with a recent 2017 reference strain (MT357898) (99.4% identity).

**Fig. 4.**
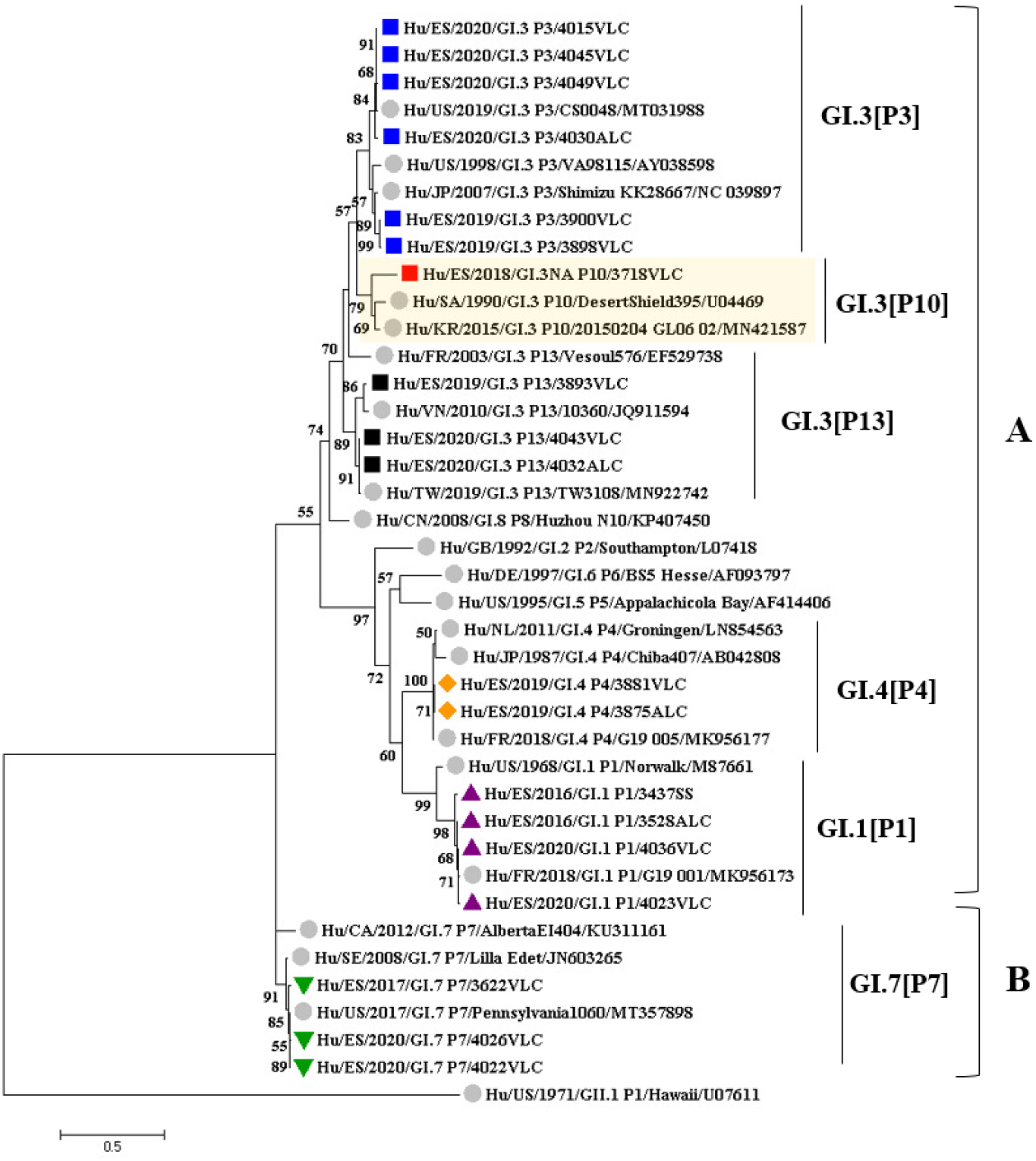
Phylogenetic analysis of the ORF1-ORF2 junction region (RNA polymerase and VP1 genes) of 19 GI norovirus collected from 2016 to 2020. The evolutionary history was inferred using the maximum likelihood method based on the Kimura 2-parameter model with a bootstrap of 1,000 replicates. Bootstrap values >50% are shown. The tree is drawn to scale; the branch lengths measure the number of substitutions per site. The analyses included 40 sequences (19 samples and 21 references). The GI.3P[10] “untypeable” strain and two reference sequences (U04469 and MN421587) are highlighted in yellow. There were 480 positions in the final dataset including nucleotides from 5157 to 5636. strains are represented by their GenBank accession numbers and indicated with grey filled circles. Sequences obtained in this study are indicated as follows: ▴ in purple, GI.1 [P1]; ▪ in blue, GI.3 [P3]; ▪ in red, GI.3 “untypeable” [P10]; ▪ in black, GI.3 [P13]; ♦ in orange, GI.4 [P4] and ▾ in green, GI.7 [P7].

##### Emergence of a novel GI.3 capsid genotype

The partial capsid sequence analysis of GI norovirus strains showed high variability in the GI.3 capsid genotype (Fig. 5A). Norovirus GI.3 strains segregated into three well defined clusters, according to the associated polymerase genotypes (P3, P10 and P13) (Fig. 5A). Intragenic variability was observed within genotypes GI.3[P3] and GI.3[P13]. There were isolates that clustered with old and new variants. For instance, the GI.3[P3] subcluster exhibited three different subgroups. One of them showed a high identity (99.1%) with a recent 2019 reference strain (MT031988), another shared a common ancestor with an older 2007 reference strain (NC_039897) and a third (“4030ALC” isolate) was phylogenetically more distant, with a nucleotide identity of between 94.5% and 97.5% compared to reference strains. On the other hand, the isolate 3718 could represent a new capsid genotype. When genotyping was performed using the Norovirus typing tool (https://www.rivmn.nl/mpf/norovirus/typingtool) no genotype was assigned to the capsid. Phylogenetic analysis showed that it clusters with Asian and African GI.3 untypeable strains (Fig. 5A). Sequence identities within this cluster ranged from 93% to 100%. This cluster has a common ancestor with GI.3[P13] strains, and surprisingly, presents greater genetic distance from the GI.3 variants [P10] such as Desert Shield (U04469) (21.1%). Furthermore, SimPlot analysis was performed using the recombinant GI.3 untypeable [P10] virus (3718) as a query sequence (Fig. 5B). Data showed that GI.3[P10] Desert Shield (U04469) is one of the parental sequences providing the polymerase region, and GI.3 untypeable [P10] (MW255366) is highly similar to the query recombinant strain 3718. However, we did not identify the parental sequence that provides the capsid region. It could be hypothesized that this parental sequence is a common ancestor from subcluster GI.3 untypeable (Fig. 5A). However, the recombination breakpoint was detected at nucleotide position 5367 in the ORF1 relating to the MT089579 genome. GI.3 untypeable capsid (1-98 aa, VP1) was further analyzed through alignment with ClustalW (Fig. 6), revealing a single amino acid change (aa 32, S-A) in the GI.3 untypeable strains. The mutation was non-synonymous, replacing a serine (uncharged amino acid) with an alanine (hydrophobic amino acid). Although the change was conservative, it is striking that the alanine in this position was conserved within the GI.1[P1] genotype.

**Fig. 5.**
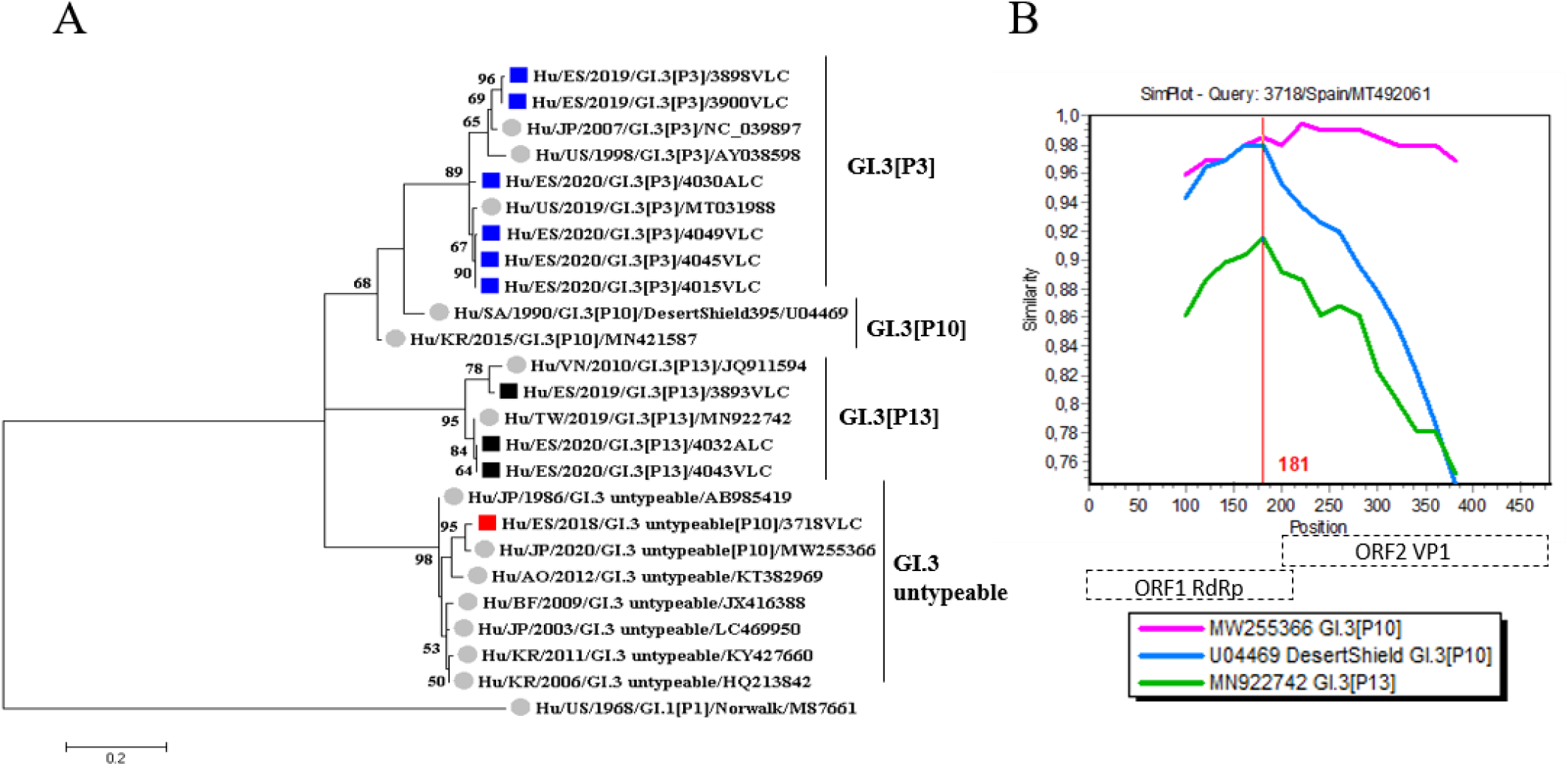
Analysis based on the partial VP1 sequence (ORF2) of GI.3 norovirus collected from 2018 to 2020. A) Phylogenetic tree of 290 bp within the capsid region (5’ ORF2). Reference strains are named after their GenBank accession numbers and indicated with circles. Sequences obtained in this study are indicated as follows: ▪ in blue, GI.3 [P3]; ▪ in red, GI.3 untypeable [P10]; ▪ in black, GI.3 [P13]. The scale bar at the bottom of the tree indicates distance. Bootstrap values (1000 replicates) are shown at the branch nodes and values above 50% are shown. B) SimPlot analysis of Spain GI.3[P10] norovirus. A SimPlot was constructed using SimPlot version 3.5.1 with a slide window width of 200 bp and a step size of 20 bp. The X-axis indicates the nucleotide positions in the multiple alignments of the norovirus sequences; the Y-axis shows percentage of similarity between norovirus reference strains and the query sequence.

**Fig. 6.**
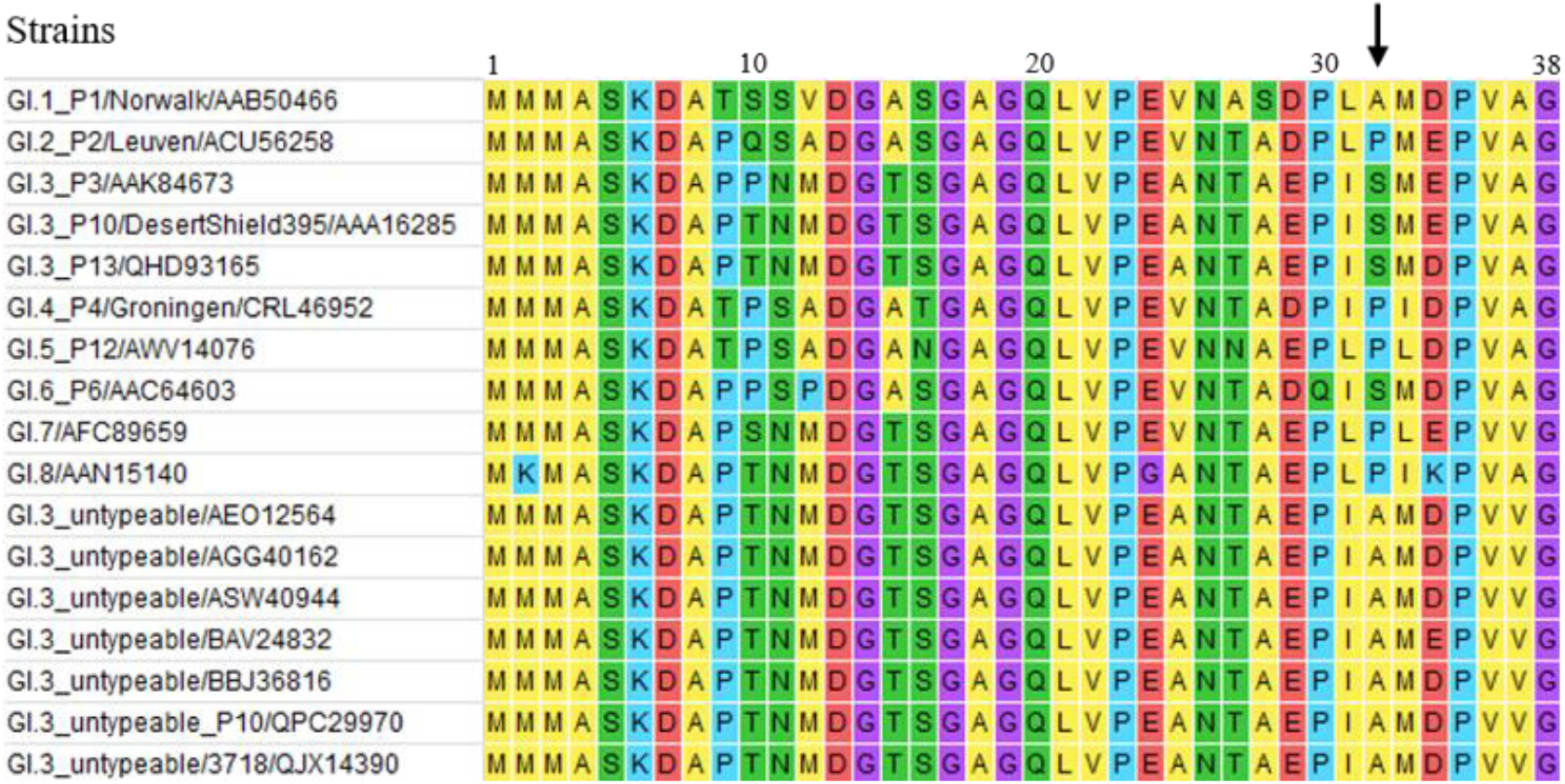
Amino acid changes within a partial VP1 sequence (1-38 aa) of a GI.3 untypeable strain compared to reference strains. A specific amino acid change in the GI.3 genotype is marked with an arrow. Colors indicate amino acid category as follows: yellow, hydrophobic; green, uncharged; blue, positively charged; red, negatively charged; purple, special.

#### Genetic analysis of norovirus genogroup II

##### GII.4 recombinants

Since 2016, three recombinant strains of the GII.4 Sydney variant have been identified as causative agents of gastroenteritis in Spain. The GII.4 Sydney[P31] and GII.4 Sydney[P4 New Orleans] strains were predominant during 2016 to 2018, but were displaced in 2019 by the recombinant GII.4 Sydney [P16] strain. There were GII.4 untypeable viruses associated with P4 New Orleans and P31 polymerase. We analyzed the genetic variability of the GII.4 strains by performing phylogenetic analysis of a partial sequence of the VP1 of 53 GII.4 noroviruses (Fig. 7). GII.4 norovirus strains were divided into four clusters, corresponding to GII.4 Sydney[P16], GII.4[P4 New Orleans], GII.4 Sydney[P31] and GII.4[P31] genotypes. GII.4 Sydney[P16] strains grouped together and shared a recent common ancestor with GII.4 untypeable[P4 New Orleans] strains and GII.4[P31]. The GII.4 Sydney[P31] lineage grouped with several reference strains and shared a close common ancestor with the GII.4[P4 New Orleans] cluster. Interestingly, GII.4 untypeable[P31] strains segregated in an independent cluster separate from the remaining groups.

**Fig. 7.**
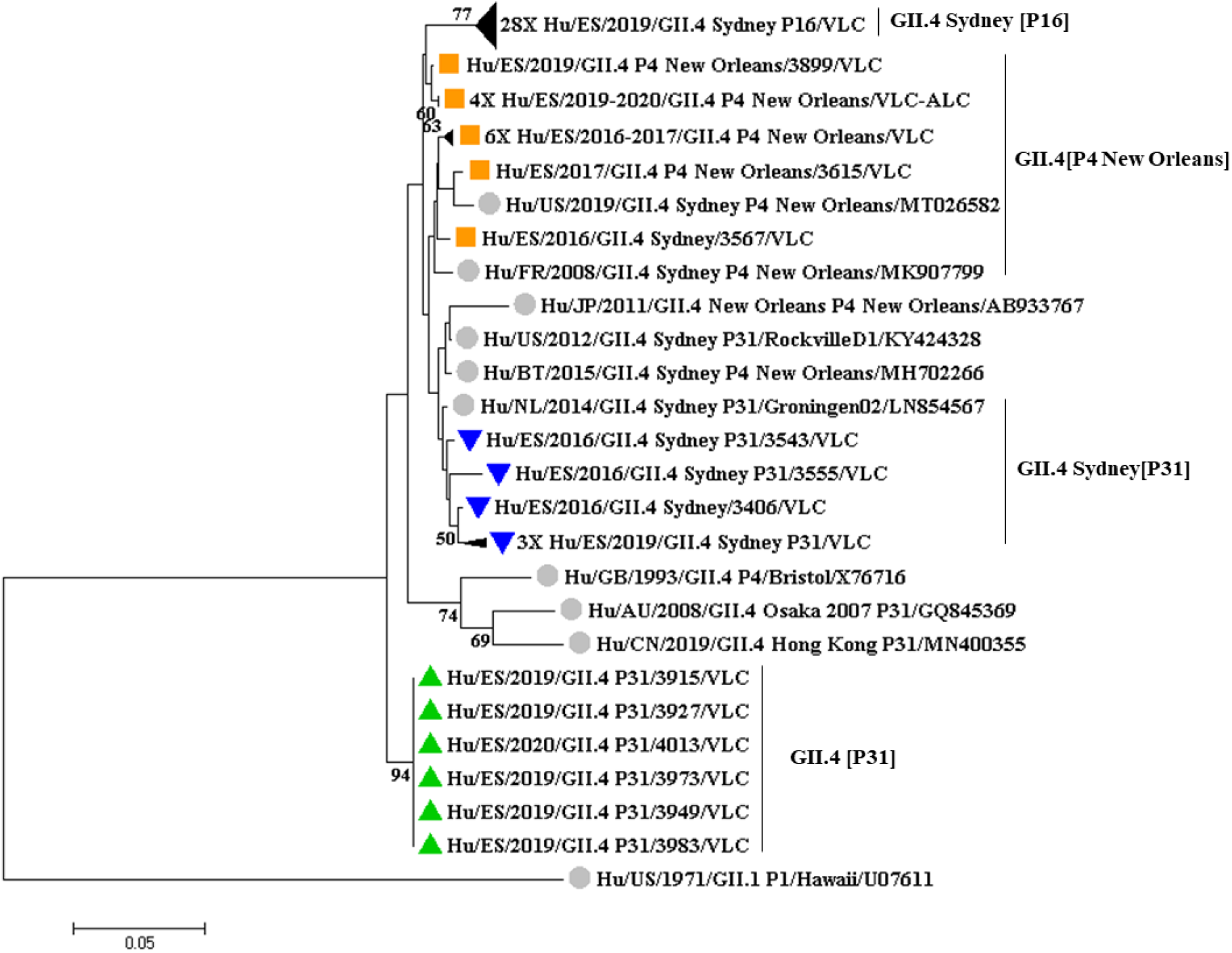
Phylogenetic analysis of norovirus GII.4 capsid collected from 2016 to 2020. The evolutionary history was inferred using the neighbor-joining method based on the Kimura 2-parameter model with a bootstrap of 1,000 replicates. Bootstrap values >50% are shown. The tree is drawn to scale; the branch lengths measure the number of substitutions per site. The analyses included 67 sequences (53 samples and 14 references). There were 276 positions in the final dataset including nucleotides from 5096 to 5372. strains are represented by their GenBank accession numbers and indicated with grey filled circles. Sequences obtained in this study are indicated as follows: ▪ in orange, GII.4 [P4 New Orleans]; ▾ in blue, GII.4 Sydney[P31]; ▴ in green, GII.4 [P31].

### Molecular phylogenetic characteristics of recombinant GII noroviruses

The recombinant norovirus GII strains characterized in this study were sorted into six well-supported clusters (A to F) in the phylogenetic tree, corresponding to 15 different recombinant genotypes (Fig. 8). SimPlot analysis showed recombination breakpoints near the ORF1/2 junction region for all strains (Fig. 9). In fact, the recombination breakpoints were identified at varying positions from nucleotides 129–301, matching the nucleotides positioned at 5011–5132 in relation to the reference viral genome NC_029646, localized in the ORF1 region for six genotypes (GII.3[P12], GII.3[P21], GII.3[P30], GII.6[P7], GII.13[P21], GII.14[P7]), in the ORF1-ORF2 overlap for GII.5[P40] and in the ORF2 region for two other strains (GII.1[P33] and GII.4 Sydney[P31]).

**Fig. 8.**
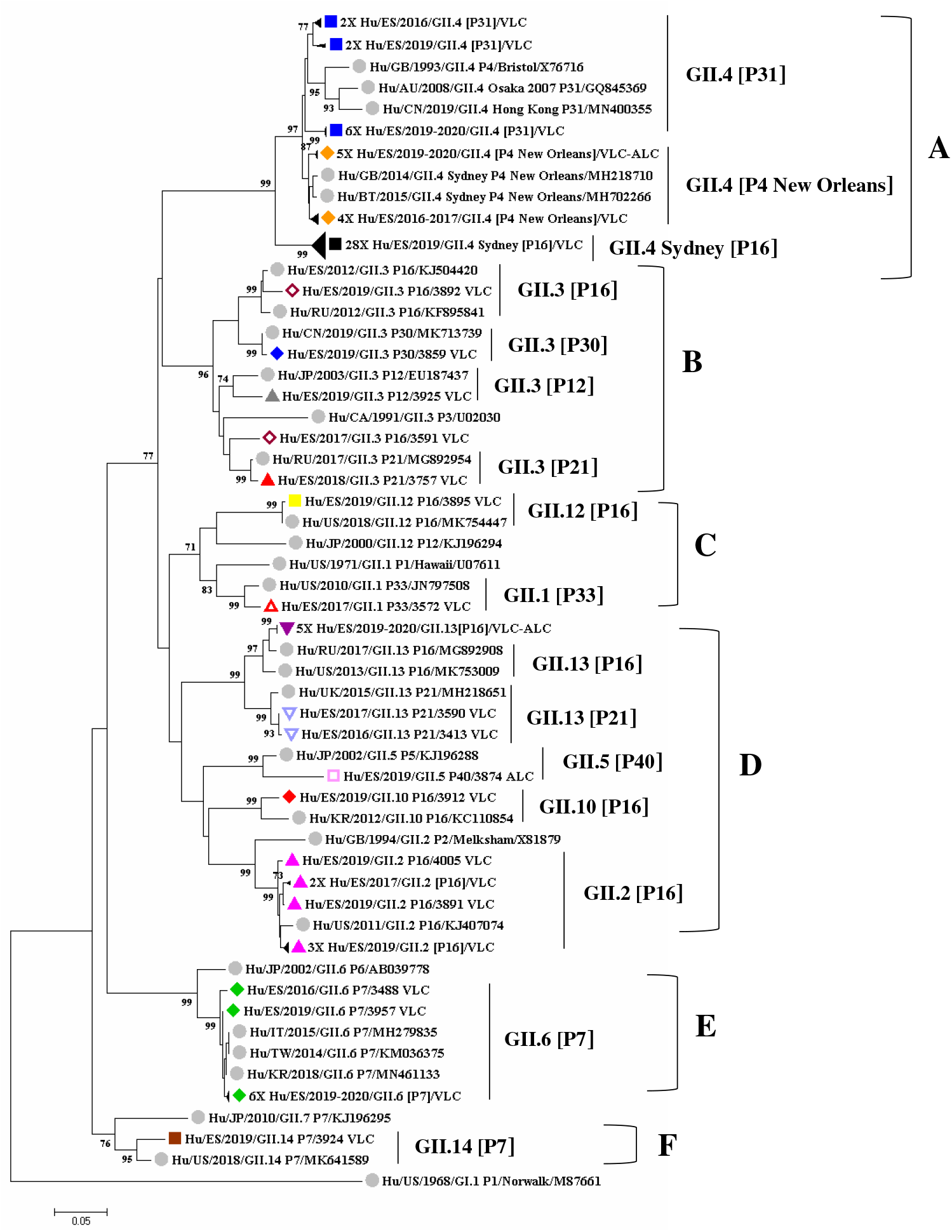
Phylogenetic analysis of recombinant noroviruses GII (ORF1-ORF2 junction region) collected from 2016 to 2020. The evolutionary history was inferred using neighbor-joining method based on the Kimura 2-parameter model with a bootstrap of 1000 replicates. Bootstrap values >70% are shown. The tree is drawn to scale; the branch lengths measure the number of substitutions per site. The analyses included 108 sequences (79 samples and 29 references). There were 417 positions in the final dataset including nucleotides from 4936 to 5352. strains are represented by their GenBank accession numbers and indicated with grey filled circles. Sequences obtained in this study are indicated as follows: ▴ empty in red, GII.1 [P33]; ▲ in pink, GII.2 [P16]; ▲ in grey, GII.3 [P12]; ♦ empty in garnet, GII.3 [P16]; ▲ in red, GII.3 [P21]; ♦ in blue, GII.3 [P30]; ♦ in orange, GII.4 [P4 New Orleans]; ▪ in black, GII.4 Sydney [P16]; ▪ in blue, GII.4 [P31]; ▪ empty in pink, GII.5 [P40]; ♦ in green, GII.6 [P7]; ♦ in red, GII.10 [P16]; ▪ in yellow, GII.12 [P16]; ▼ in purple, GII.13 [P16]; ▾ empty in blue, GII.13 [P21] and ▪ in brown, GII.14 [P7].

**Fig. 9.**
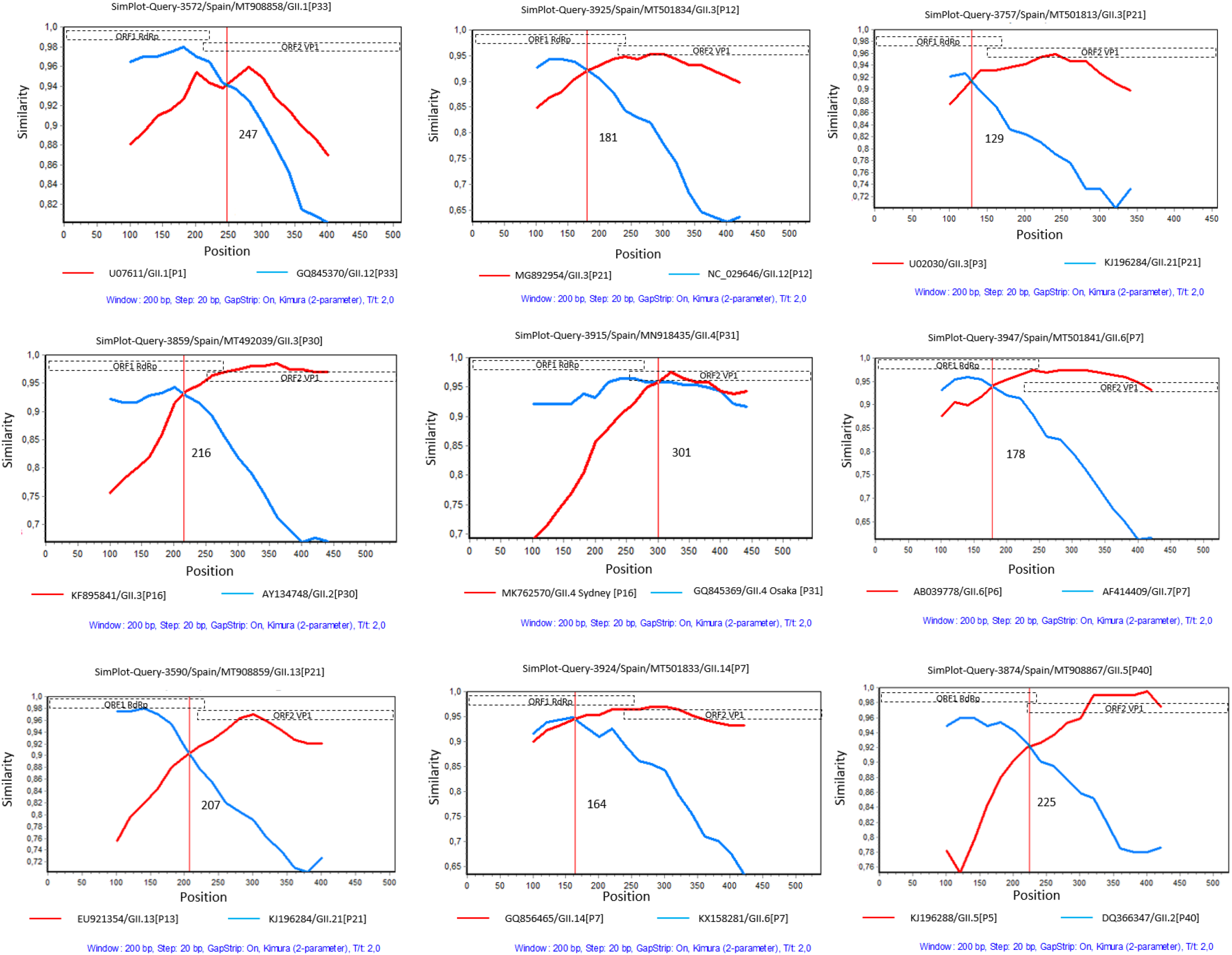
SimPlot analysis of nine norovirus recombinant strains sequences. Graphs were constructed using SimPlot software version 3.5.1 with a slide window width of 200 bp and step size of 20 bp. At each position of the window, the query sequence was compared to each of the reference strains. The X-axis indicates the nucleotide positions in the multiple alignments of the norovirus sequences; and the Y-axis shows the percentage of similarity between norovirus reference strains and the query sequence. The Kimura 2-parameter model is applied.

## DISCUSSION

Recombination is among the most important genetic diversity-generating events within the *Norovirus* genus (18). The current study provides an overview of the molecular epidemiology of noroviruses drawn from analysis of clinical samples collected in the Valencian Community, Spain, from January 2016 to April 2020. GII was the predominant genogroup (78.1%), with GII.4 Sydney 2012 found to be the most prevalent genotype followed by GII.2, GII.3, GII.6 and GII.17, which concurs with globally circulating strains (36, 37).

Herein, we report norovirus infections caused by 18 different recombinant strains for the first time in Spain, including uncommon genotypes such as GI.5[P4] and GII.5[P40]. A high prevalence of recombinant strains (90.4%) was observed among genogroup GII viruses from 2018 to 2020. In contrast, only five recombinant strains were detected in a previous study carried out in 2009–2012 in the Basque Country (Spain) (38). Remarkably, most outbreaks during the 2019–2020 period were caused by GII.4 Sydney strains associated with P16 and P31 polymerases. The emergence of these strains coincided with a high prevalence of GII.2[P16] viruses in different Asian and European countries (39, 40). Note that emergence of these genotypes did not lead to major amino acid changes in the capsid of GII.4 (41, 42), suggesting that the evolutionary advantage lies in the polymerase. Interestingly, in this study we found GII.P16 polymerase harboring six different capsids (GII.2, GII.3, GII.4 Sydney, GII.10, GII.12 and GII.13). However, this ‘promiscuity’ was not unique to the P16 polymerase, as P21 was also detected combining with GII.3 and GII.13. We also found GI.3 and GII.3 viruses associated with different polymerases (P10 and P13) and (P12, P21 and P30), respectively. The versatility of multiple polymerases combining with different capsids supports the idea that the polymerase plays a crucial role in norovirus infection. Furthermore, a recent evolutionary analysis of 25 different polymerases pinpointed high evolutionary ratios between P4, P12, P16 and P31 strains, suggesting that the RdRp region evolves rapidly, similarly to VP1 (43). A meta-analysis of norovirus infections recently carried out in China reported that recombinant strains carrying the GII.P16 RdRp genotype such as GII.2 and GII.4 have caused epidemics, and concluded that the GII.P16 polymerase may trigger pandemics (44).

In the present study we identified a new subcluster of GII.4 Sydney associated with the P31 polymerase, which clusters into an outgroup in the phylogenetic analysis of the VP1 region of GII.4 (Fig. 7). This new subcluster of GII.4 Sydney has also recently been reported in Chile, the United States, Germany, Australia and New Zealand, albeit associated with the P4 polymerase (36). Further analysis of the ORF2 of these isolates is warranted to examine possible amino acid changes in antigenic epitopes of VP1, which could allow the virus to evade the immune response. A new GII.4 variant could emerge if strains from this subcluster continue evolving and spread worldwide, accumulating capsid changes. A recent study has shown that pandemic GII.4 variants require pre-adaptation to the host, circulating in the population for up to nine years before emerging globally (45). This further supports the importance of epidemiological surveillance of norovirus infections.

In genogroup I, GI.3 and GI.4 viruses were the most common genotypes in this study, in accordance with previous reports (46, 47). The presence of recombinant strains is lower in this genogroup than in GII, although we observed both GI.3[P13] and GI.3[P10], as well as the more uncommon GI.5[P4]. Widespread circulation of GI.3[P13] strains has been reported in Taiwan (48). Nevertheless, there is little awareness of emerging new variants among GI viruses. Herein, we have identified an isolate (strain 3718) which could represent a new genotype, or at least a new variant of the GI.3 capsid. Indeed, when the genotype was determined using the Norovirus typing tool, no genotype was assigned for the capsid, which was classified as P10 for the polymerase gene. Furthermore, this isolate clustered together with unassigned Asian and African strains to form an outgroup, based on phylogenetic analysis of a partial capsid region with GI.3 reference strains (Fig. 5). According to the recently redefined norovirus classification criteria, a strain belongs to a genotype if it has >85% sequence identity in VP1 (16). In this case, strain 3718 has a genetic distance of more than 20% (1–98 aa region of VP1) compared to reference strains GI.3 [P10] and GI.3 [P13]. In addition, a single amino acid change (aa 32, S-A) was detected in the GI.3 unassigned strains. Therefore, complete ORF2 sequences of the strains from this cluster are needed to analyze possible changes in the antigenic region of the capsid, which could help avoid antibody neutralization. It is worth noting that although GI.3 is a static genotype (25), it was recently shown to have a large number of sites under positive pressure, which could explain the emergence of new variants (49). For this reason, GI infection surveillance should be encouraged by establishing classification criteria for new variants within genotypes such as GI.3.

In summary, this study reports a diverse range of recombinant norovirus strains circulating in Spain in 2016–2020. A total of 18 recombinant strains have been detected, an increase that may be due to improved genotyping procedures such as dual-typing assays. These changes have highlighted the need to unify detection criteria for noroviruses targeting the ORF1-ORF2 overlapping region, in order to characterize the genetic diversity of recombinant noroviruses worldwide.

## ACKNOWLEDGMENTS

This study was partially supported by a grant from the Spanish Ministry of Science and Innovation, Carlos III Health Institute (grant PI16/01471) and by a research grant to NN-L from the Conselleria d’Educació, Cultura i Esports, Generalitat Valenciana (grant ACIF/2020/076).

